# Low temperature triggers sociality in a facultatively social bee

**DOI:** 10.1101/2025.10.03.680320

**Authors:** Jan J. Kreider, Matt C. Elmer, Harley Thompson, Marina Pizarro-Krkljus, Vanessa Kellermann, Ido Pen

**Author notes:** Equally contributing authors (Vanessa Kellermann, Ido Pen).

## Abstract

The evolutionary transition to eusociality in the solitary ancestors of social insects, like ants, termites, wasps, and bees, may have been initiated by an environmental induction of group living. Plasticity in social behaviour in response to the environment can be found in facultatively social insects. However, it is currently unknown which environmental cues trigger the individual decision of an insect to nest solitarily or socially. Such environmental cues could be identified by exposing individuals of a facultatively social insect to experimentally manipulated cues in a controlled environment. Here, we report such an experiment with the facultatively social allodapine bee *Exoneura robusta*, which we exposed to different temperature and predation treatments in climate chambers. We observed that low temperature triggered the bees to behave more socially. Furthermore, there was substantial variation in the tendency to be social among individual bees. Our study identifies a specific environmental cue that triggers a plastic response in sociality. Such plasticity could be crucial at the origin of group living and social breeding, facilitating the evolution of eusociality in insects.

## Introduction

Eusocial insects – ants, termites, some wasps, and some bees – have evolved a complex form of sociality where typically one reproductive queen is supported by thousands of sterile workers [1]. Discovering how eusociality has evolved has been a core objective of evolutionary biology for a long time [2–5]. In 1987, West-Eberhard proposed that sociality could initially be induced by a change in environmental conditions, rather than by genetic mutation [6]. This would lead to the emergence of facultative sociality, where individuals plastically adjust their social behaviour depending on the environment; thus, under certain environmental conditions social nesting would be favoured over solitary nesting and vice versa [7,8]. Such behavioural plasticity has been demonstrated in a sweat bee, where solitary bees from northern populations become social when transplanted to the south and social bees from the south become solitary in the north [9,10]. This suggests a role of climate or a correlate of climate (e.g., season length [11,12]) in causing sociality, but the trigger for the individual decision to nest solitarily or socially is unknown.

Facultative sociality can be found in several species of bees [8] and wasps [13]. Within bees, facultative sociality is most well-known from the halictine bees (e.g., species of the genera *Halictus* [14], *Lasioglossum* [15], and *Megalopta* [16]). The decision of individual bees to engage in solitary or social nesting depends on the environmental conditions in some species [9,10,17], but in others there is a genetic underpinning for solitary vs. social behaviour [18–20]. A second major group of bees with facultatively social species are the xylocopine bees, e.g., the carpenter bees [21], the small carpenter bees [22], and the allodapine bees [23]. Although it is well-known that sociality can be induced by the environment in many facultatively social species, it remains elusive which environmental cues trigger sociality.

*Exoneura robusta* (Allodapini) is an Australian facultatively social bee, which can nest solitarily and socially in the same population [24,25]. *E. robusta* progressively provisions its brood in an undivided nest tunnel, at the end of which the larvae feed on a pollen ball [24]. Females disperse in spring and establish a nest either solitarily or together with other females [26,27]. Newly co-founded nests are typically initiated by females that have previously shared a nest and which therefore are related [24,26,27]. In such newly co-founded nests, most females breed and only a low level of ovarian differentiation and reproductive skew exist among the nest members [24,28]. Females, who do not disperse, reuse their natal nest. In these reused nests, one female monopolizes reproduction in late winter/spring while her nestmates assume subordinate helper roles [29–34]. This is achieved by pheromonal inhibition of ovarian development in the subordinate females [32] and by aggressive behaviour towards mated subordinate females that are excluded from the nest by the dominant female, who guards the nest entrance [31,35]. Behavioural specialisation occurs in autumn with some subordinate females specialising in foraging and feeding the individuals in the nest by trophallaxis, others building the nest tunnel, and some are inactive [36]. Since reproductive dominance is determined by the order of adult eclosion [29,30], a monopolisation of early reproduction allows the dominant female to pass on reproductive dominance to her own offspring [31]. A secondary reproductive may lay some eggs later during the season, but many females never reproduce [28,37]. During the breeding season, early emerging females can provide some care towards the developing larvae in the nest [33]. Populations of *E. robusta* differ in the frequency of co-founding of new nests [38] (and the possibility for early emerging females to help rear developing larvae [23,39–41]), but it is unclear which cues from the external or internal environment of an individual trigger the bee to be in a solitary or social nest. Such environmental cues could be identified under highly controlled conditions where specific cues are manipulated. However, only few attempts to maintain facultatively social insects in the laboratory have been successful [42,43]. To our knowledge, the only study that has manipulated environmental cues has shown that the availability of nesting opportunities in *E. robusta* has no effect on the tendency of individuals to engage in solitary vs. social nesting, demonstrating that sociality is not caused by nest site limitation [38]. Therefore, there is a need for an experiment targeting the manipulation of specific environmental cues with the aim of altering the proportion of solitary vs. social nesting in a facultatively social insect.

Apart from cues from the external environment, a behaviourally plastic response in social behaviour could also be triggered by individual properties, such as relative body size of an insect. Coercion of smaller daughters into helping has been demonstrated to underlie reproductive division of labour in small carpenter bees [44,45]. This is potentially because small body size limits an individual’s reproductive ability [5], which should favour helping behaviour in small individuals. On the other hand, in groups of wasps [46,47], body size often determines the outcome of agonistic interactions and thereby the position of individuals in the reproductive hierarchy, which should favour social grouping behaviour in large individuals that become reproductively dominant. Consequently, an experiment targeting the manipulation of environmental cues should ideally also investigate if body size affects individual tendencies towards solitary or social nesting.

Here, we investigate if temperature or brood predator cues are involved in the decision of a bee to nest socially or solitary in the facultatively social bee *E. robusta*. We selected temperature as a candidate cue for inducing sociality because sociality may be energetically advantageous over solitary nesting at lower temperature [48,49]. We chose olfactory predator cues because group-living *Exoneura* have better defence against brood predation by ants; single females of solitary nests leave their nests unguarded during foraging, while social nests always have guards due to behavioural specialisation and division of labour between the nestmates [24,25,35,36,50]. Furthermore, we investigate whether body size predisposed the bees towards solitary or social behaviour.

## Materials & Methods

### Field work

We sampled nests of *Exoneura robusta* on 17 April 2023 along Lyrebird Gully Creek and Rifle Range Gully in the Dandenong Ranges National Park, Victoria, Australia (Fig. S1) from *Cyathea australis* tree fern fronds, in which *E. robusta* nests (Fig. 1a-c). We chilled nests over night at 4 °C at Monash University before opening them. In autumn, nests of *E. robusta* contain adult females and males that emerged during the summer along with a small number of females that emerged the previous year and survived the breeding season. In total, 139 nests contained 553 *E. robusta* females, 38 *E. robusta* males, 80 *Inquilina schwarzi* (a social parasite bee of *E. robusta*) females, and 39 *Inquilina schwarzi* males (Fig. S2). Within the study population, during the breeding season of 2023 approximately 64% nests were found to be social. Average temperature ranges from 8.1 °C to 20.6 °C, average daily minimum from 5.0 °C to 15.1 °C, and average daily maximum from 11.3 °C to 27.2 °C at the study site (each averaged over one month; years 2013-2018; Fig. S3)

**Figure 1.**
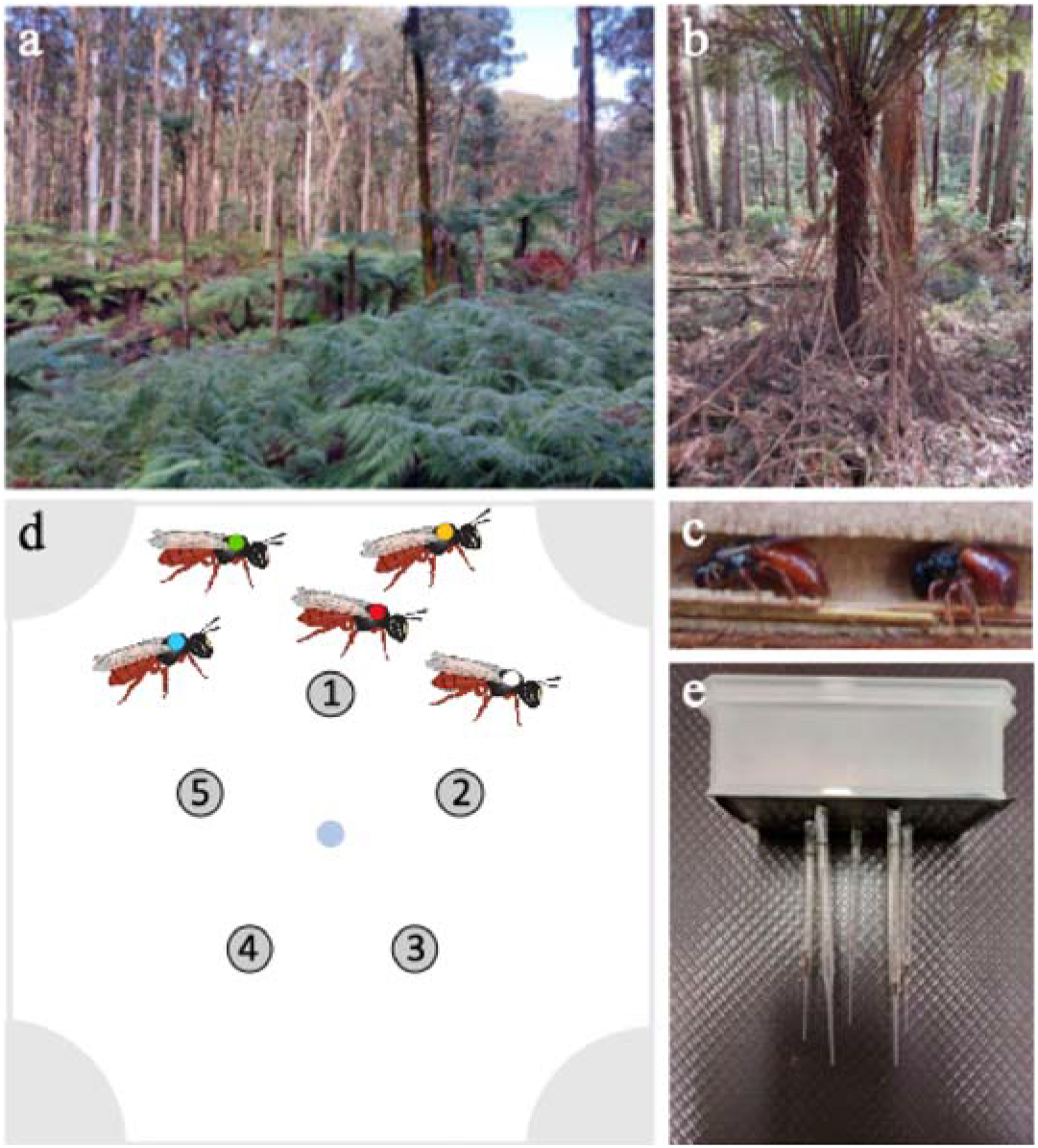
Natural habitat and nesting substrate of *Exoneura robusta* and experimental setup. **(a)** *E. robusta* occurs in forests along the Australian east coast. **(b)** The tree fern *Cyathea australis* has pithy stems in which *E. robusta* excavates a tunnel as a nest. **(c)** Nest tunnel in longitudinally-opened tree fern front with two female *E. robusta* bees. **(d)** In our experiments, five individually-marked bees from five different nests were released into an experimental enclosure with five nesting opportunities (circles with numbers). Food was provided in the middle of the enclosure (blue dot). A predator cue or control cue was provided in the four edges of the enclosure. **(e)** Enclosures consisted of plastic boxes with Pasteur pipettes glued onto them to imitate nests. The Pasteur pipettes were covered by a cardboard box such that the nest tunnels were dark.

### Behavioural experiment

We individually colour-marked all *E. robusta* female bees (water-based Posca pen). In the experiment, we only used female bees that had recently emerged during the summer and had not had the chance to disperse from their natal nests. We achieved this by excluding females with wing wear which indicates that they had foraged during the previous breeding season and survived until the end of the season, i.e., these females are old and emerged the previous season. We set up 72 groups of five female bees each (360 bees in total). Natural nests of *E. robusta* most often consist of one to four bees during the breeding season [24]. Therefore, the number of bees in a nest was not unnaturally restricted by our experimental setup with five bees per group. The bees within each group had previously not shared a nest and were therefore unfamiliar with each other, thus preventing recognition by former nestmates [27,51]. Each group was housed in a separate enclosure with five artificial nests (Pasteur pipettes; Fig. 1d+e, Fig. S4), allowing the bees the choice to nest solitarily or socially with others. The bees occupied the bottom of the Pasteur pipettes, accumulating closely to each other with physical contact, as they do in natural nests. They spent most of the time in the artificial nests and only left those for foraging, showing that Pasteur pipettes can conveniently be used for experimental setups and observation with *E. robusta*, complementing other observation nests [43]. We divided the 72 groups across six treatments with three different temperatures (18 °C, 22 °C, 26 °C) and the presence/absence of an olfactory predator cue of ants that occur near natural nests of *E. robusta* (12 replicates per treatment; details in Fig. S4). To mimic a predator cue, we obtained ant nests and foraging ants from the field site close to the nests of *E. robusta*. We crushed one individual of *Polyrhachis femorata*, two individuals of *Colobopsis gasseri* and three individuals of *Myrmecorhynchus emeryi* onto Whatman filter papers. We added a fresh filter paper to renew the scent cue on each day of the experiment. We let the bees acclimatize in their treatment for 2h before opening the entrances to the artificial nests. We recorded the locations of all individually-marked bees over five days once every 2h during the 12h daily activity period (lights in the climate chambers were on from 7:00am to 7:00pm, i.e., 12:12 light cycle). This resulted in a total of 30 observations per individual bee over the five days. In the centre of each enclosure, we provided fresh food every day (1:1 sugar water) in a capillary tube, such that bees had unlimited food access. We froze all bees at the end of the experiment and shipped them to the University of Groningen on dry ice, where they were stored in a -70°C freezer for further analyses. We obtained measures of wing length for all 360 bees from the experiment. Wing length of the left and right forewing were measured under an Olympus SZ51 stereo microscope with an eyepiece graticule from the base of the wing to the merging of the marginal cell with the front side of the wing under 15x magnification (Fig. S5). Wing length is commonly used as a proxy for body size [24,29].

### Statistical analysis

We performed all statistical analyses with R 4.2.2 [52], using the Bayesian statistics package *brms* [53–55] in combination with the MCMC sampler of *cmdstanr* [56]. We considered a bee to be “social” at a particular time when there was at least one other bee in the nest with her at that time, whereas a bee was considered “solitary” if she was in the nest alone. To model how the proportion of bees in different nest types (solitary vs. social vs. outside nest) varied over time and between treatments, we used a Markov chain model that encapsulates the transition probabilities between nest types. These probabilities were allowed to vary between treatments, to depend on individual body size (i.e., wing length) and to change over time. Random effects of individual ID, replicate, and individual nest-of-origin were included. We performed pairwise comparisons between treatments by computing the posterior odds ratio for each comparison and assessed significance by evaluating the overlap of the 95% credible interval with unity. To evaluate the significance of the random effects, we ran models with and without them and computed their difference in the leave-one-out (LOO) cross validation information criterion. We estimated how much of the observed variance could be attributed to the various random and fixed effects by calculating the variance over the posterior probability of social nesting for each random effect and the fixed effects. Further details on the statistical analyses can be found in the Supplementary Materials.

## Results

The probability of bees to nest socially increased with decreasing temperature (Fig. 2a). In the absence of predator cues, the bees were more likely to nest socially at 18 °C and 22 °C than at 26 °C (Fig. 2b). Similarly, in the presence of predator cues, the bees were more likely to nest socially at 18 °C compared to 22 °C and 26 °C, but not more likely social at 22 °C compared to 26 °C (Fig. 2b). This suggests an interaction between temperature and predator cues. However overall, predator cues did not induce social nesting behaviour (Fig. 2c). Furthermore, body size played no role in the tendency of bees to nest socially (mean: -0.02 [95% CI: -0.10, 0.06]; pd = 0.64). Individual bees had different tendencies to be social. The variation explained by the nest-of-origin in the natural population and the between-replicate variation was lower than the variation explained by between-individual differences (Fig. 3, Table 1).

**Table 1.** Leave-one-out information criteria (LOOIC) for the full model corresponding to Fig. 2 and models that exclude one of the random effects each. A lower LOOIC indicates a better model.

| LOOIC | SE | ΔLOOIC | ΔSE | Model |
| --- | --- | --- | --- | --- |
| 10321.7 | 146.4 | 0.0 | 0.0 | full |
| 10323.9 | 146.3 | 2.3 | 4.2 | excl. nest-of-origin |
| 10480.6 | 147.6 | 158.9 | 23.7 | excl. individual ID |
| 10329.9 | 146.6 | 8.2 | 5.9 | excl. group (replicate) |

**Figure 2.**
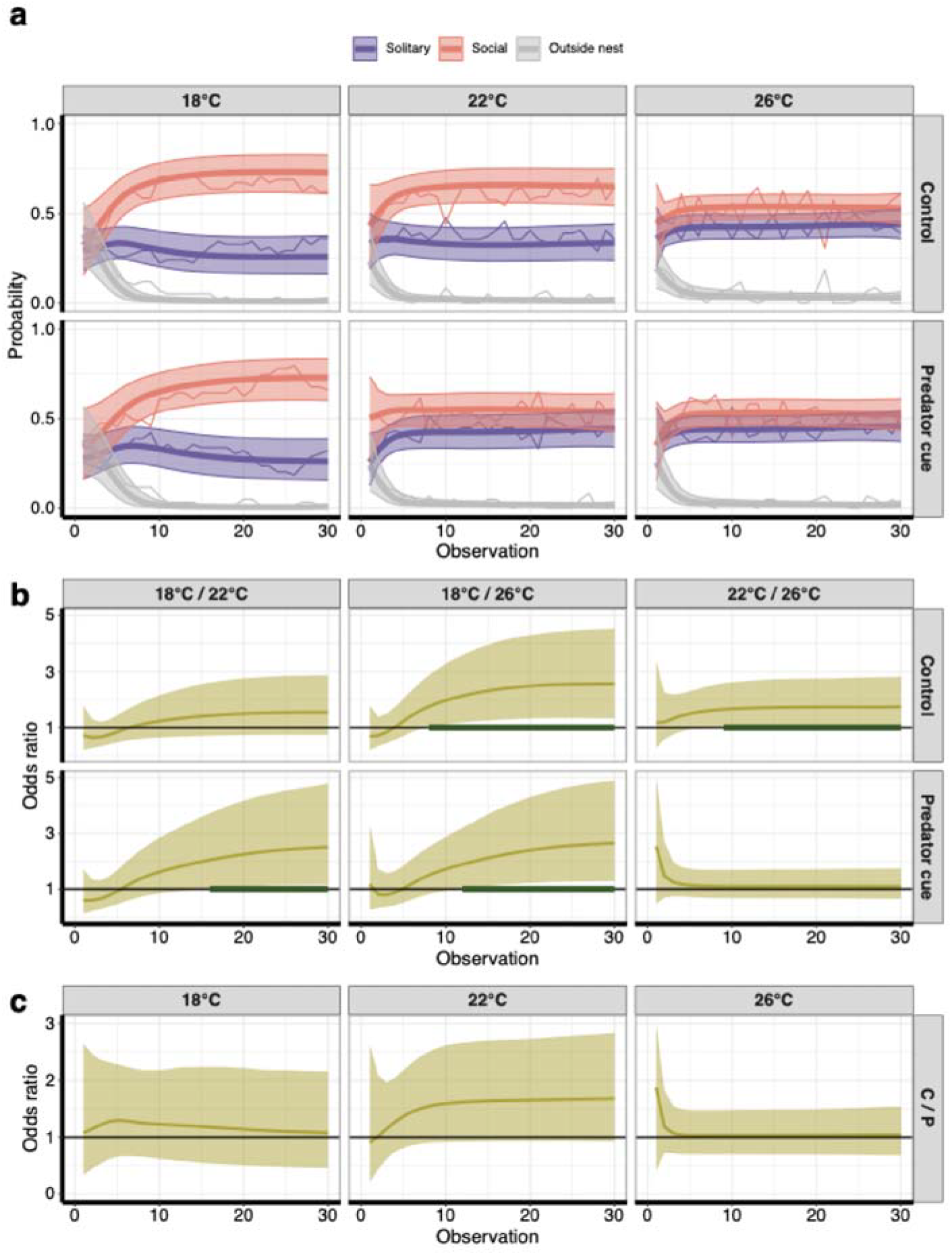
Behaviourally plastic response in sociality to temperature and a brood predator cue in *Exoneura robusta*. **(a)** Probability of bees to be solitary (purple), social (red) or outside the nest (grey). Time represents 30 observation moments. The bold line represents the posterior mean and the area around it the 95% credible interval of the mean from a time- and treatment-dependent Markov chain model. The thin lines connect the empirical means calculated from the raw data. **(b)** Odds ratios for sociality vs. solitary or outside between the temperature treatments for the control group and the predation treatment. **(c)** Odds ratios for sociality vs. solitary or outside for the comparison between control (C) group and the predation (P) treatment at the same temperature. **(b + c)** The thick lines represent the posterior mean odds ratios and the shaded area around it the corresponding 95% credible interval (CI). A comparison can be considered statistically significant when the 95% CI does not overlap with unity (black line). Time points at which a comparison was statistically significant are marked with a thick green horizontal bar.

**Figure 3.**
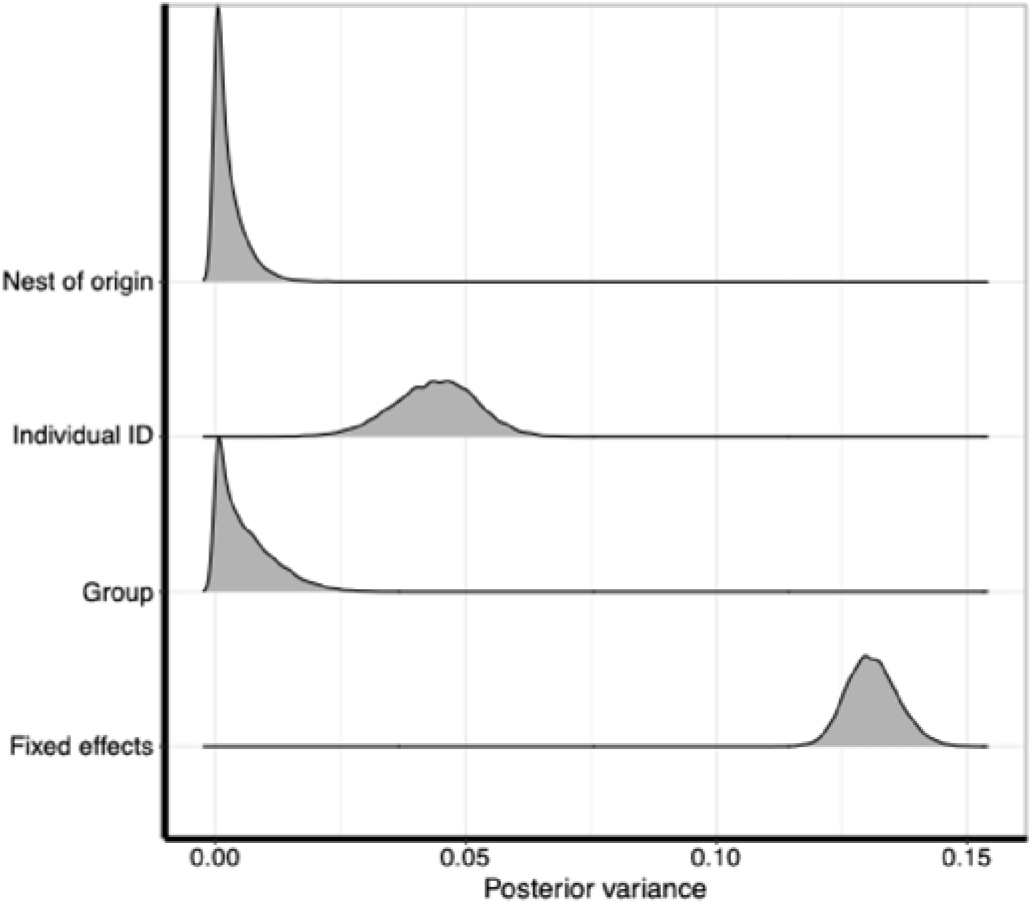
Posterior variance of the random effects (nest-of-origin, individual ID, and group/replicate) and the fixed effects (temperature, predation treatment, and body size).

We observed that there were differences in nest switching between treatments (note that this refers to which *nest* the bees were in but not which *nest type*, i.e., social vs. solitary vs. outside nest). Nest switching occurred at higher frequencies with increasing temperature (all three pairwise comparisons pd = 1.0; Fig. 4) and was more likely in the control compared to the predation treatment at 18°C (pd = 0.99) and vice versa at 22°C (pd = 0.97). There was no difference at 26°C (pd = 0.71). Body size had no effect on the nest switching tendency (−0.05 [-0.14, 0.04]; pd = 0.80). Some variation originated from interindividual behavioural differences, i.e., individual differences in tendency to switch nests, but little variation in nest switching tendency could be attributed to the nest-of-origin and replicate (Table S1).

**Figure 4.**
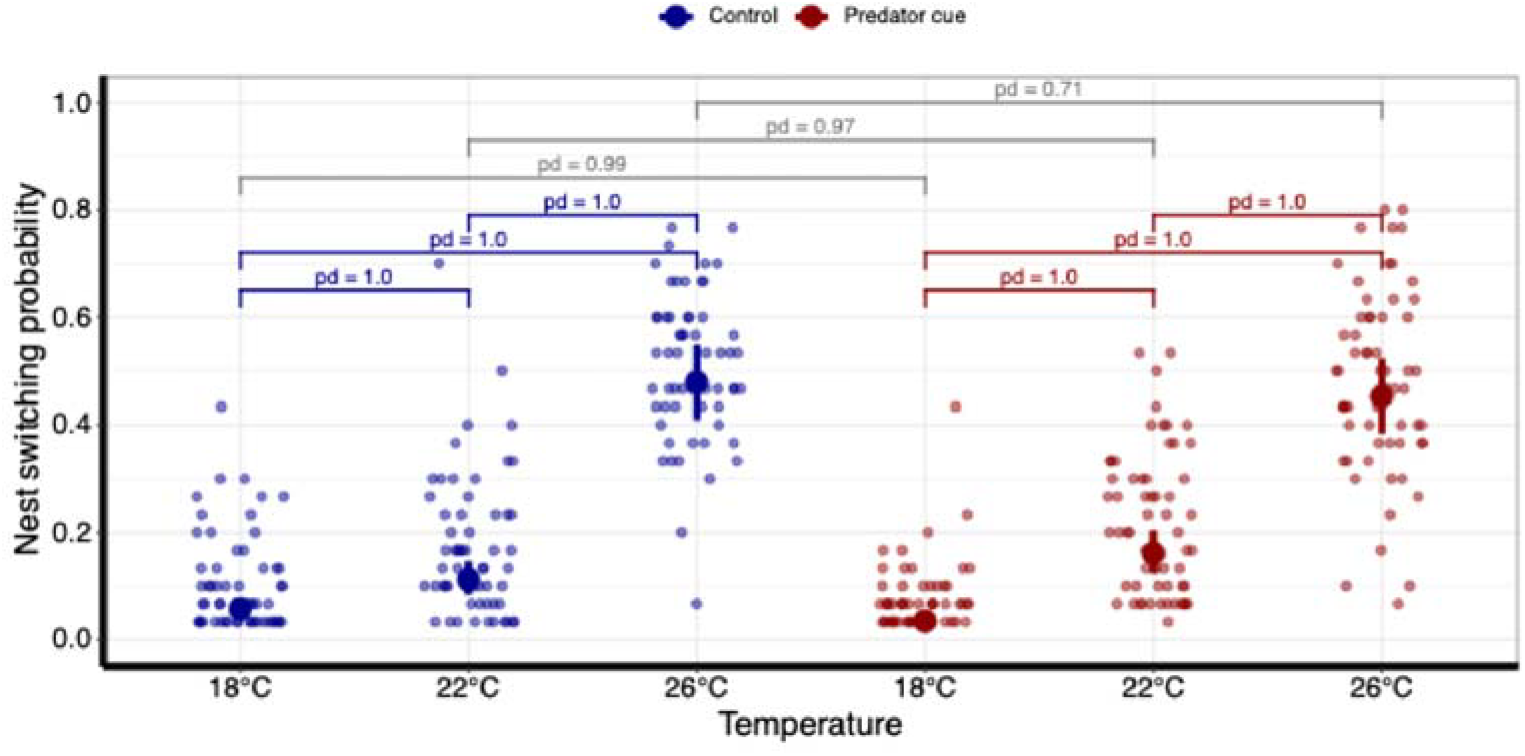
Nest switching varies by temperature and predation treatment. Probability of bees to switch between nests across temperatures in the control (blue) and predator cue treatment (red). Large dots represent the posterior mean and error bars the 95% credible interval of the mean. Small dots are the nest switching probability of an individual bee throughout the experiment.

To test if these differences in nest switching between treatments could have biased our result that low temperature triggers social nesting, we performed simulations in which we investigated the effect of nest switching tendency on the probability of social nesting. We assumed five equally-spaced nests between which five individuals switch at random. We then varied the switching probability and recorded the proportion of social nests. The proportion of bees in social nests was not influenced by differences in the nest switching probability (Fig. S6), demonstrating that the differences in nest switching at different temperatures (Fig. 4) were not underpinning our result that temperature influenced the probability to nest socially (Fig. 2).

## Discussion

Here, we report an experimental induction of sociality in a facultatively social insect by manipulation of a specific environmental cue. Geographic variation in social behaviour can be found across social arthropods [11,57], including *Exoneura robusta* [23,38–41]. However, to date no study has experimentally identified specific environmental cues that cause changes in social nesting. In our experiment, low temperature triggered an increase in the frequency of social nesting. This plastic response to temperature in *E. robusta* is the reverse of that in sweat bees [8], which are more likely social at lower latitudes [9,10,58,59], where temperatures are typically higher. In sweat bees, warmer environments allow the production of two broods per season and sociality emerges when the individuals from the first brood become workers and help raise the second brood. In *E. robusta*, social nests occur throughout the winter and spring [24], during which the bees might experience thermal benefits from group living [48]. In the population examined, *E. robusta* frequently experiences low temperatures between 5 °C and 10 °C during winter and spring (Fig. S3). Therefore, one possible reason we see a preference for social nesting at cooler temperatures is the potential for thermal benefits. It has been shown in carpenter bees, that group-living bees more successfully conserve heat and body mass than individuals in solitary nests [48]. Further research should measure the metabolic effects of sociality in *E. robusta*. Field studies should investigate whether a correlation of sociality with temperatures occurs in natural populations of *E. robusta*.

In contrast to the effect of temperature on sociality, we observed no overall effect of predator cues on the decision of bees to nest socially or solitarily. We chose an olfactory cue of different ant species (mixing multiple species increased the chance of at least having one brood predator) that occur near the natural nests, as these species are likely predators of *E. robusta* brood [25]. Formic acid has been shown to elicit behavioural responses in honeybees [60] and olfactory predator cues of ants have been given in filter papers in other behavioural experiments [61]. Although this predator cue did not have an overall effect on social behaviour, we still cannot exclude the possibility that other predator cues would elicit a response in social behaviour. It could also be that the predator cue we used would show an effect on behaviour at a different stage in the *Exoneura* life cycle when brood is present. Future research should evaluate behavioural responses to other predator cues, e.g., to living ants, spiders, or crickets, and at different colony life cycle stages.

Our experiment has been conducted in autumn with recently emerged female bees that have not been through a natural dispersal period, similar to the dispersal experiment by Schwarz and Blows [27]. In autumn, bees in natural colonies become behaviourally specialised since specialisation in guarding the nest entrance allows the dominant female to exclude other mated females from the nest [31,35]. In our behavioural experiment, the bees were therefore given a choice to group or occupy a nest solitarily just before the establishment of behavioural roles and development of ovarian size differences, which leads to reproductive division of labour. We found rather strong differences in the individual tendency of bees to be in a social or solitary nest. One reason for these individual differences could have been body size differences between individuals, where either smaller or larger bees have an increased probability to engage in social nesting. However, we found no effect of body size on social behaviour. Since we conducted the experiment prior to the development of ovarian size differences between individuals [32], it is also unlikely that large differences in ovarian size underlie the individual differences. It is known that *E. robusta* can recognize familiar females from their nest and are more likely to re-group with those in dispersal experiments [26,27]. Maintaining associations with familiar females enables co-founding of new nests with kin. In our experiment, we used females that had not previously shared a nest to exclude association between familiar individuals. Therefore, the bees within an enclosure were on average unrelated. Since some individuals of *E. robusta* mate in autumn before ovarian maturation [31], there could have been differences in mating status between individuals, which could perhaps explain the interindividual differences to some extent. Since the firstly emerging female in natural nests mates and then excludes other females to obtain reproductive dominance, mating status should also correlate with age (time since emergence) [29,30]. Being accepted into a social nest could depend on the mating status of the female and the mating status of the other females in the enclosure. If more females were mated, this should lead to more rejection of nestmates and fewer social nests. As a consequence, we should observe variation between replicates caused by random variation in mating status of the five bees, which also was the case. However, whether this underpins interindividual differences in social behaviour requires further investigation.

We observed that individuals sometimes switched between nests with differences in nest switching frequencies between treatments, which was a consequence of higher activity levels at higher temperatures. Although nest switching occurred, the proportion of bees in solitary vs. social nests stabilised over time and we showed in simulations that nest switching had no effect on the probability of observing bees grouped in a nest. Nest membership in *E. robusta* is reported to be rather stable [24], but nest switching occurs in other social insects where individuals become social parasites that lay eggs in colonies of non-relatives [62]. There is also one record of nest switching and even nest fusion in *E. robusta* from observation nests [36], but a quantification of nest switching frequency in natural populations has not been performed.

We employed a definition of sociality where bees were considered social when they grouped with other bees in the same nest [63,64]. This grouping behaviour indeed seems to involve social interaction, since the bees accumulated closely to each other at the bottom of the artificial nests with physical contact. Since we conducted the experiment in autumn before the establishment of behavioural roles and reproductive differentiation, the bees would presumably develop behavioural specialisation. However, breeding in captivity is not yet feasible in this species, prohibiting a continuation of our experiment. Group formation and the subsequent emergence of behavioural specialisation and division of labour are key steps during the evolutionary transition from solitary to eusocial breeding. It has repeatedly been found that sociality in some bees [8] and wasps [13] is plastic. Group formation could have initially been a plastic response to the environment [6]. Alloparental care behaviour and division of labour could have emerged from pre-existing regulatory responses, not requiring their *de novo* evolution [65–67], e.g., derived from maternal care behaviour that is heterochronously expressed towards a nestmate’s offspring [68,69]. Thus, key steps during the transition from solitary to eusocial insects – group living and alloparental care – could have originated through plasticity and exaptation of existing regulatory responses [70–72], which could have further been adapted by natural selection (genetic accommodation [7,73,74]) and ultimately evolved to be expressed irrespective of the environment (genetic assimilation [7,75]), yielding irreversibly committed queen and worker castes.

Lastly, behavioural traits, such as social behaviour, are often highly flexible to changes in environmental conditions. Yet, we have a poor understanding of how changing environments impact bees and their social behaviour [76]. Our results suggest that sociality in *E. robusta* may be altered by climate change, such that increasing temperatures will likely reduce the probability of social nesting in this species – opposite to the effect on sweat bees [77]. However, several environmental cues could affect social behaviour in natural populations and interact in intricate ways. Understanding the environmental drivers of sociality is needed to predict the consequences of climate change on bee behaviour and the flow-on effects on pollination services and ecosystem functioning.

## Author contributions

JJK, VK, and IP conceptualized the research. JJK, MCE, and HT did the field work. JJK, MCE, HT, MPK, and VK conducted the experiments and lab work. JJK and IP analysed the data. All authors contributed to the writing of the manuscript.

## Acknowledgements

We are grateful to Mike Schwarz for providing his expertise on *Exoneura robusta* to us. We thank Kim Holzmann for designing the bee in Figure 1. We would like to thank the Center for Information Technology of the University of Groningen for providing access to the Hábrók high performance computing cluster. This research was supported by a Travelling Fellowship by The Company of Biologists, a Dobberke grant by the Royal Netherlands Academy of Arts and Sciences, an Academy Ecology Fund grant by the Royal Netherlands Academy of Arts and Sciences, and a Student Research Grant by the Animal Behavior Society, all to JJK. Furthermore, JJK was supported by an Adaptive Life grant by the University of Groningen.

