## Supplement for "Low temperature triggers sociality in a facultatively social bee"

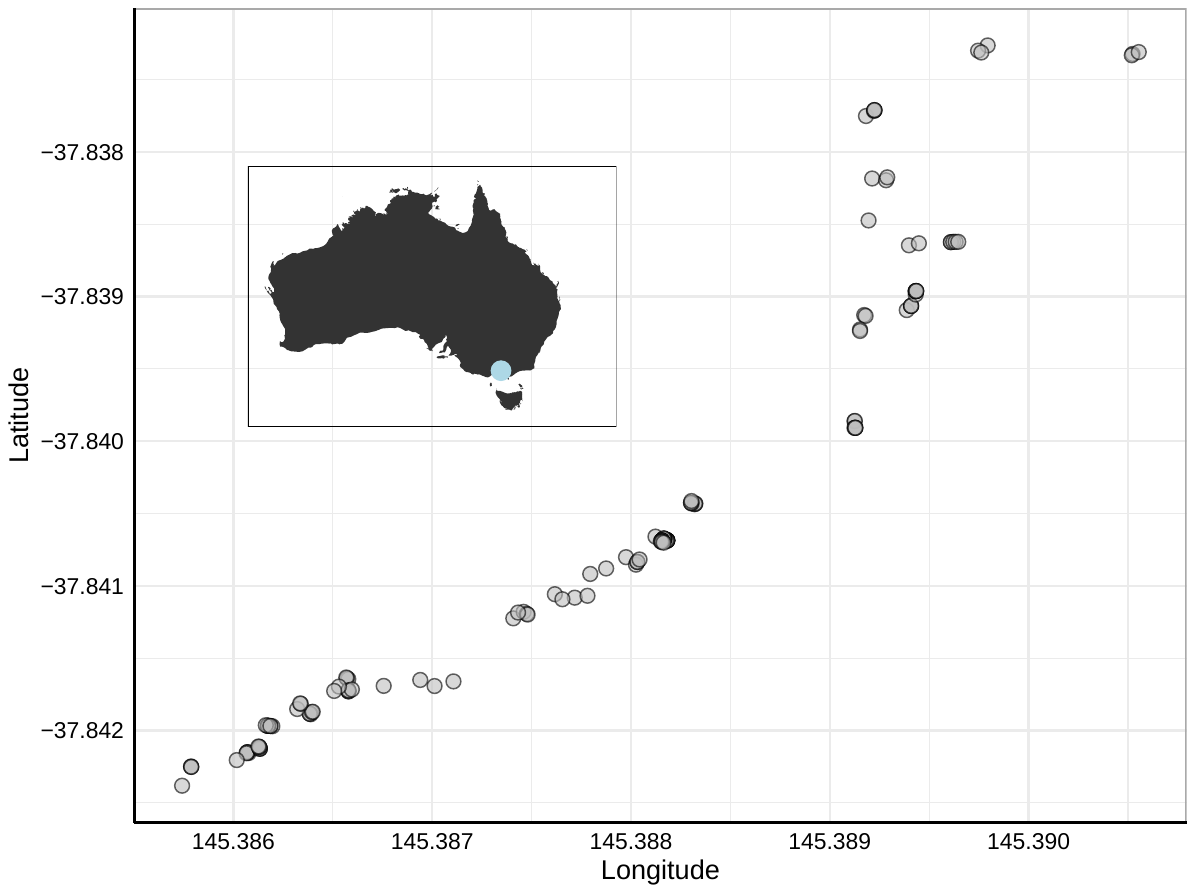

**Fig. S1** Sampling locations of *Exoneura robusta* nests along Lyrebird Gully Creek and Rifle Range Gully in the Dandenong Ranges National Park, Victoria, Australia (light blue dot on map left top). Each grey dot represents the sampling location of a nest.

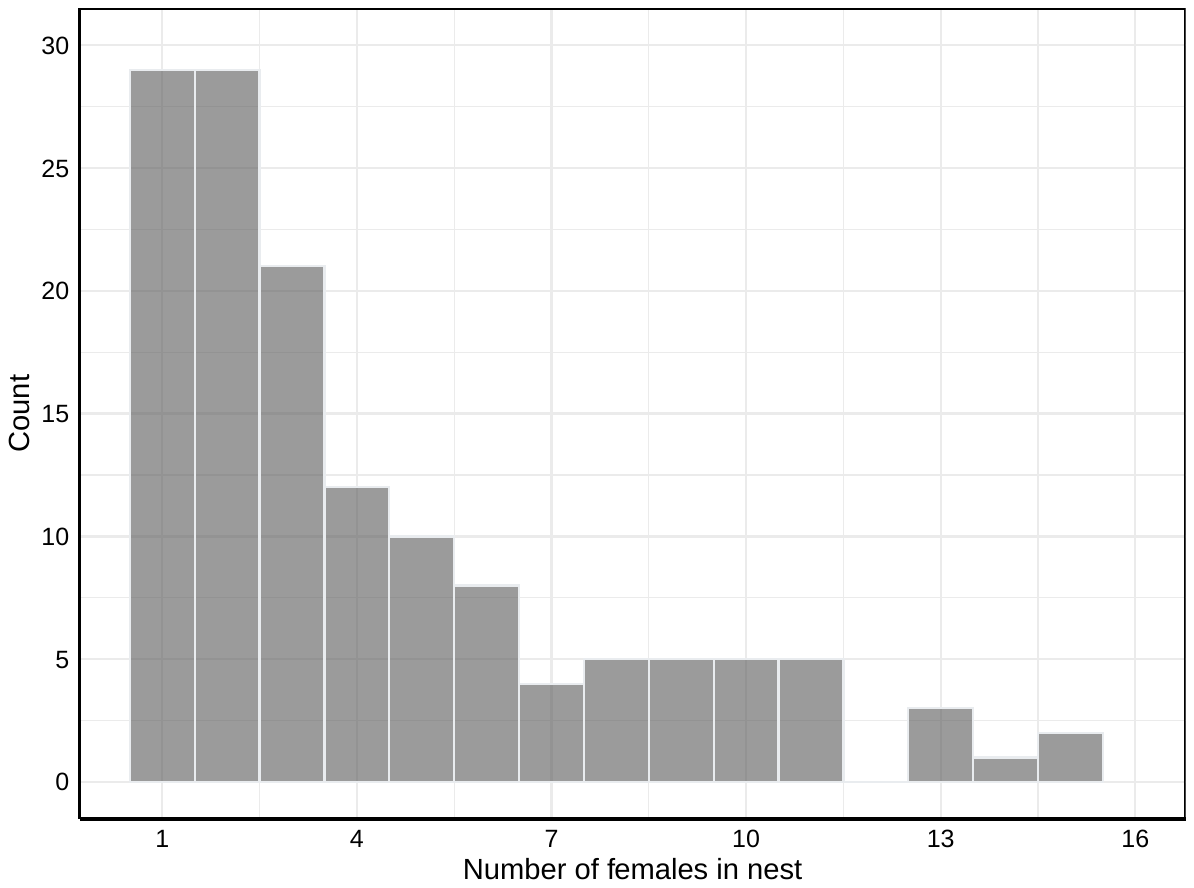

**Fig. S2** Histogram of number of females per nest in natural populations in autumn. Number of females represents the number of *Exoneura robusta* and *Inquilina schwarzi* females combined.

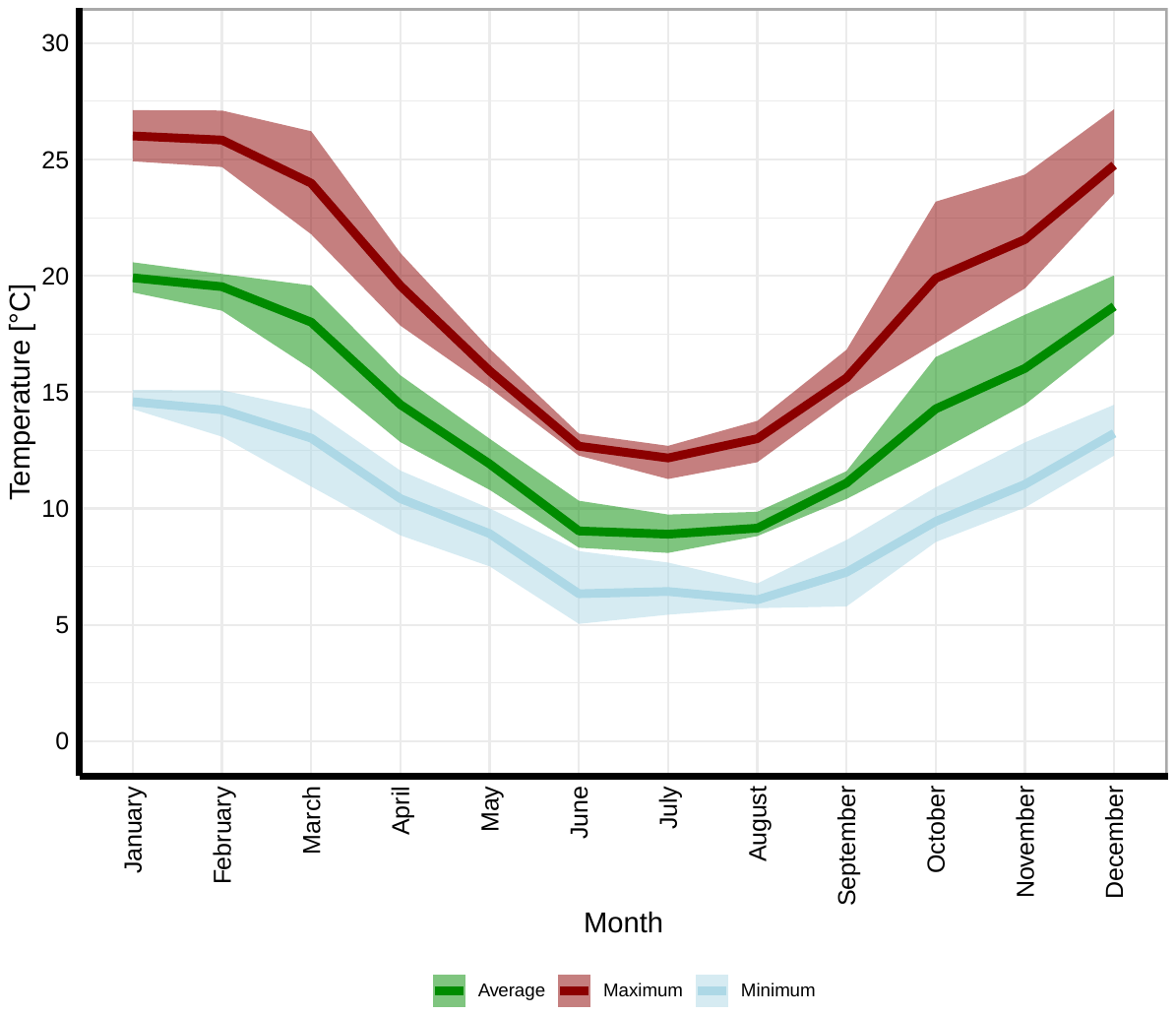

**Fig. S3** Average, average daily maximum, and average daily minimum temperature at the field site over the years 2013-2018^1^. The line represents the mean and the ribbon the range across the years.

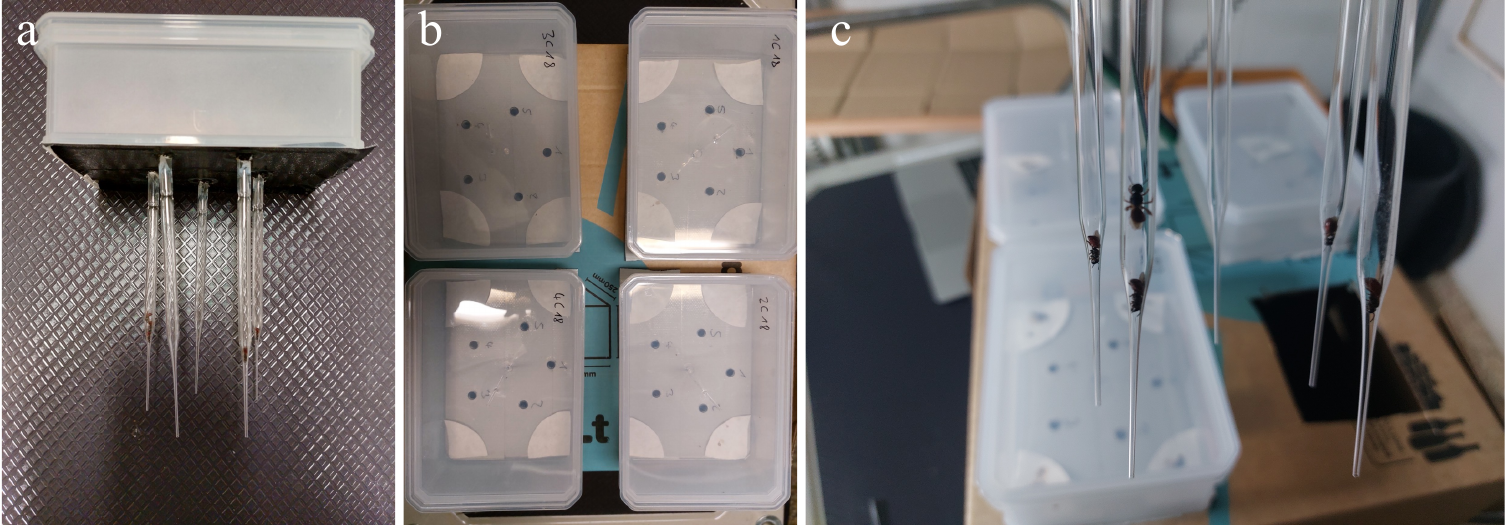

**Fig. S4** Enclosures used for the experiment with *Exoneura robusta*. **(a)** Each enclosure consisted of a plastic box with five artificial nests (Pasteur pipettes). **(b)** The enclosures were positioned on a cardboard box so that the artificial nests were dark. The edges of the enclosures had Whatman filter papers with ants squeezed into them or empty. **(c)** The bees settled quickly in the artificial nests. The Pasteur pipettes allow for easy monitoring of the nest of individuals at every data recording moment.

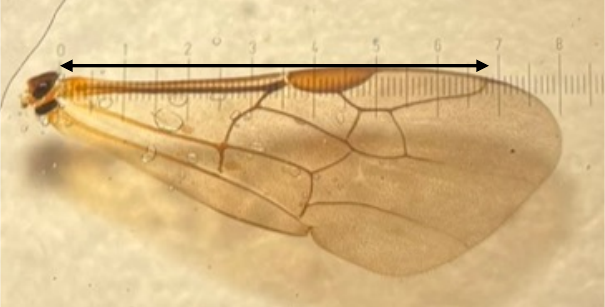

**Fig. S5** A wing of *Exoneura robusta*. The arrow shows the measurement of the wing length from the base of the wing to the merging of the marginal cell with the front side of the wing.

**Table S1.** Leave-one-out information criteria (LOOIC) for the full model corresponding to Fig. 4 and models that exclude one of the random effects each. A lower LOOIC indicates a better model.

| LOOIC | SE | ∆LOOIC | ∆SE | Model |
| --- | --- | --- | --- | --- |
| 8839.8 | 109.8 | 0.0 | 0.0 | full |
| 8838.9 | 109.7 | -0.9 | 0.5 | excl. nest-of-origin |
| 9144.7 | 111.0 | 304.8 | 33.6 | excl. individual ID |
| 8841.0 | 109.6 | 1.2 | 2.9 | excl. group (replicate) |

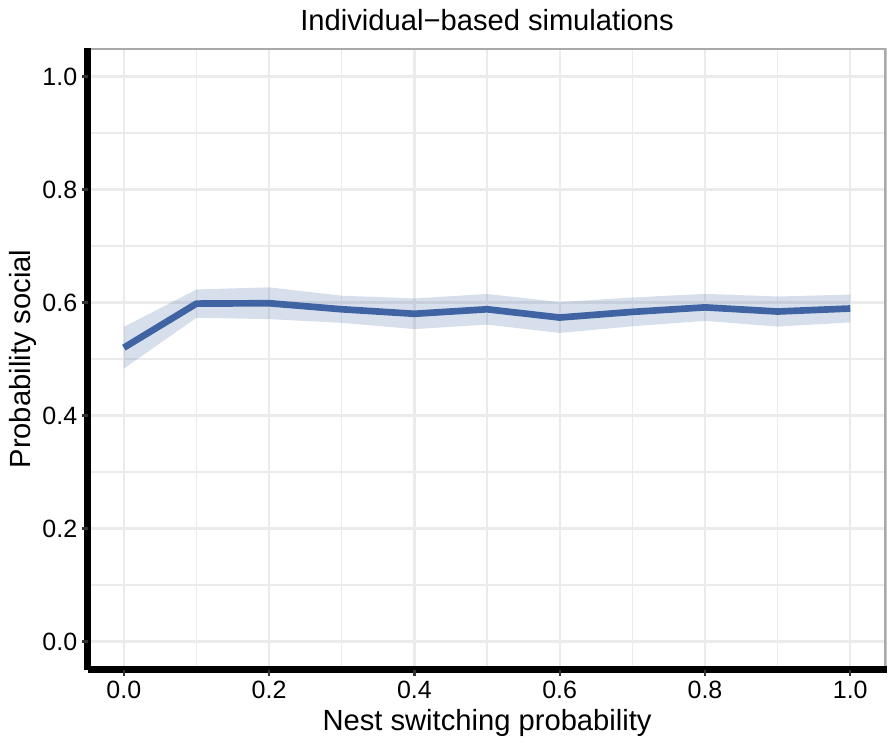

**Fig. S6** Probability of individuals to be in a social nest by different nest switching probabilities in individual-based simulations. The blue line represents the mean and the area around it the 95% confidence interval of the mean. We simulated each nest switching probability between 0.0 and 1.0 in steps of 0.1 with 10 replicate simulations for each parameter setting.

**Details on the Bayesian model corresponding to Figure 2**

We modelled the transition probabilities between the three states (solitary, social, outside nest) as a Markov process in discrete time^2^. Accordingly, a bee in state $Z_{t}=x\in\{A = \mathrm{solitary}, B = \mathrm{social}, C = outside nest\}$ at time *t* transitions to state $Z_{t+1}=y$ at *t* + 1 with probability

$\Pr(Z_{t+1}=y|Z_{t}=x)\equiv p_{x\to y}$	(S1)

We can write a transition matrix with the transitions between states *A*, *B* and *C*

$\mathbf{M}=\left[ \begin{matrix} p_{A\to A} & p_{B\to A} & p_{C\to A} \\ p_{A\to B} & p_{B\to B} & p_{C\to B} \\ p_{A\to C} & p_{B\to C} & p_{C\to C} \end{matrix} \right]$	(S2)

Since the columns of **M** sum to 1, **M** has six independent entries. We allowed **M** to vary by treatment $T\in\{18C, 18P,22C,22P,26C,26P\}$, where *C* refers to control and *P* to predator cue, standardized body size *S*, and the observation moment *t*, where this temporal effect was modelled by a thin-plate spline function^3–6^ that varied by the previous state $Z_{t-1}$. Furthermore, we added a unique replicate identifier, a unique individual identifier, and a unique nest-of-origin identifier as random effects. We modelled the initial distribution of the bees into the three states by the same predictors and random effects, and estimated correlations between random effects for the initial distribution and the transition probabilities.

We assessed alternative models without a temporal effect and with a temporal effect that did not only vary by the previous state but by an interaction of previous state and treatment. We compared models using the leave-one-out (LOO) information criterion, which showed that both alternative models performed worse than the model above.

Based on the transition probabilities $p_{x\to y}$, the probability $\Pr(Z_{t}=y)$ that a bee is in a particular state *y* at time *t* can be derived from the law of total probability

$\Pr\left( Z_{t}=y \right)=\sum_{x} \Pr\left( Z_{t-1}=x \right)p_{x\to y}$	(S3)

where the sum is over all possible states. We implemented the model with R 4.2.2^7^ in the Bayesian statistics package *brms*^3–5^ in combination with the MCMC sampler of *cmdstanr*^8^. We modelled the transition probabilities in each column of **M** by treating the observed transitions as drawn from a multinomial distribution (“categorical” response in *brms*). For logistic models with random effects at the logit scale, predictions at the probability scale are biased towards the extremes due to the nonlinearity of the logit link function. We performed bias corrections using the following formula. Given an uncorrected predicted probability $\hat{p}$, the corrected prediction is

$\hat{p}_{\mathrm{corrected}}=\hat{p}+\frac{1}{2}\sigma_{\mathrm{tot}}^{2}\hat{p}(1-\hat{p})(1-2\hat{p})$	(S4)

where $\sigma_{\mathrm{tot}}^{2}=\sum_{j} \sigma_{j}^{2}$ is the total variance of all random effects^9^.

**Nest switching probabilities**

To model the probability of nest switching in the different experimental treatments, we used logistic regression models with a binary response. Experimental treatment and body size were used as predictors, along with random effects of replicate, individual ID, and nest-of-origin. To test hypotheses, we calculated the probability of direction (pd) for each effect of interest, which represents the posterior probability that an effect occurs in a particular direction, using the “hypothesis”-function in *brms*.

**Priors, iterations, convergence and posterior predictive checks**

For all models, we used weakly informative Gaussian priors^10,11^ with mean = 0 and SD = 1 for slope regression coefficients. We used the default priors of *brms* for intercepts. The random effect parameters used the default priors of *brms* (half-t density with df = 3 for standard deviations; LKJ density for correlations). We ran the models with four chains each and discarded the first 1000 warm-up iterations, followed by 3000 sampling iterations, resulting in 12000 posterior samples. We monitored proper mixing of chains with trace plots and convergence of chains by verifying that all $\hat{R}=1.0$0. We used the *pp_check*-function from the *brms* package to evaluate model fit by inspecting posterior predictive checks.
